# Impaired Spatial Reorientation in the 3xTg-AD Mouse Model of Alzheimer’s Disease

**DOI:** 10.1101/258616

**Authors:** Alina C. Stimmell, David Baglietto-Vargas, Shawn C. Moseley, Valérie Lapointe, Lauren M. Thompson, Frank M. LaFerla, Bruce L. McNaughton, Aaron A. Wilber

**Affiliations:** Department of Psychology, Program in Neuroscience, Florida State University, Tallahassee, Florida; Neurobiology and Behavior, University of California Irvine, Irvine, California; Department of Neuroscience, University of Lethbridge, Lethbridge, Alberta, Canada

**Keywords:** Alzheimer’s disease, spatial reorientation, spatial navigation, learning and memory, sex differences

## Abstract

In early Alzheimer’s disease (AD) spatial navigation is impaired; however, the precise cause of this impairment is unclear. Recent evidence suggests that getting lost in new surroundings is one of the first impairments to emerge in AD. It is possible that getting lost in new surroundings represents a failure to use distal cues to get oriented in space. Therefore, we set out to look for impaired use of distal cues for spatial orientation in a mouse model of amyloidosis (3xTg-AD). To do this, we trained mice to shuttle to the end of a track and back to an enclosed start box to receive a water reward. Then, mice were trained to stop in an unmarked reward zone to receive a brain stimulation reward. The time required to remain in the zone for a reward was increased across training, and the track was positioned in a random start location for each trial. We found that 6-month female, but not male, 3xTg-AD mice were impaired. Male and female mice had only intracellular pathology and male mice had less pathology, particularly in the dorsal hippocampus. Thus, AD may cause spatial disorientation as a result of impaired use of landmarks.

## Introduction

Alzheimer’s disease (AD) is a progressive neurodegenerative disorder affecting over 35 million people in the world and is the leading cause of dementia (Braak & Braak 1991, Forner et al 2017, Mitchell et al 2002). AD involves the formation of amyloid beta (Aβ) plaques, followed by tau pathology in the form of neurofibrillary tangles (Braak & Braak 1991, Mitchell et al 2002). A critical problem for developing treatments that prevent AD symptomology is that Aβ and tau pathology progress substantially before serious cognitive symptoms appear, leading to a diagnosis long after neurological damage has been done (McDade & Bateman 2017). Individuals with AD have multiple cognitive symptoms including memory and navigational impairments (Henderson et al 1989, Weintraub et al 2012). In fact, one of the earliest cognitive symptoms to arise in AD is getting lost, particularly in new surroundings (Allison et al 2016, Henderson et al 1989, Weintraub et al 2012).

Essentially every mouse model designed to mimic the amyloidosis associated with AD is impaired at spatial navigation (e.g., Attar et al 2013, Liu et al 2013, Marlatt et al 2013; for review, Webster et al 2014). Impaired spatial coding, which could underlie these navigational impairments, has also been reported with less stable spatial maps in 3xTg-AD mice compared to non-transgenic (non-Tg) mice (Cayzac et al 2015, Mably et al 2017). However, the precise cause of the navigational deficit and impaired spatial coding is unclear. Some studies suggest that navigation is the primary impairment (Cañete et al 2015, Davis et al 2017, Stover et al 2015), while others suggest the impairments may actually be a consequence of failure to retain information that was learned the previous day (i.e., a memory impairment; Billings et al 2005), or a combination of memory and navigation impairments (Wood et al 2016). Further, the navigational tasks used to assess impairments all assume that mice are using distal cues to navigate the environment, but use of distal cues is not explicitly assessed.

Navigational studies in AD have often used tasks designed to engage allocentric (map-like) navigational strategies; however, there is also evidence for egocentric navigational impairments in AD. Specifically, AD can be differentiated from frontotemporal dementia (FTD) based on the presence of an additional egocentric deficit, while allocentric (map-like) impairments are present in both FTD and also AD patients (Tu et al 2017). Consistent with these findings, the parietal cortex (PC), which is thought to play a role in egocentric (e.g., viewer dependent) navigational strategies (Clark et al 2018, Nitz 2009, Nitz 2006, Nitz 2012, Whitlock et al 2012, Wilber et al 2014, Wilber et al 2017), is also dysfunctional in mouse models and humans with AD (Khan et al 2014). This dysfunction in PC was attributed to changes in a PC-hippocampal network. Emerging evidence from modeling and experimental studies suggests that this same PC-hippocampal brain network may translate the viewer-dependent (egocentric) representations of landmarks into allocentric (world-centered) coordinates for navigation (Clark et al 2018, Deshmukh & Knierim 2013, McNaughton et al 1995, Wilber et al 2014).

The ability to update knowledge of current location using surrounding landmarks would be critical for navigating new surroundings, which is dysfunctional in AD (Allison et al 2016). Here, we test the hypothesis that AD may specifically affect the ability to use distal cues to get oriented in space. In order to address this question, we selected a *spatial reorientation task* that was previously used to show deficits in aged rats and link this navigational impairment to impaired updating of the hippocampal place field map (Rosenzweig et al 2003).

We use a well-characterized triple transgenic mouse model (APPSwe, PS1M146V, and tauP301L) that expresses three major genes associated with familial AD. The 3xTg-AD mouse exhibits plaque and tangle pathologies with a distribution pattern comparable to that observed in humans (Mesulam 1999). In humans with AD, loss of synaptic density precedes neuronal degeneration (DeKosky & Scheff 1990) and is a better predictor of memory loss than plaques and tangles (Flood & Coleman 1990, Masliah et al 2001). The 3xTg-AD mouse model also mimics these synaptic changes (Oddo et al 2003).

## Methods

3xTg-AD male and female mice, and age-matched non-transgenic controls (n=5/sex), were grouped housed (2-4/cage) in 12:12 h light/dark cycles until the beginning of the experiment. Both 3xTg-AD mice (originally obtained directly from Dr. LaFerla), and non-transgenic controls of the same background strain as 3xTg-AD mice, were bred in our vivarium. We assessed male and female mice early in disease progression: 6-months (n=5/sex/genotype; N=20). All experimental procedures were carried out in accordance with the NIH Guide for the Care and Use of Laboratory Animals and approved by the University of California, Irvine and Florida State University Animal Care and Use Committees.

### Pre-training

Mice were water deprived to no less than 80% of their beginning body weight and given food *ad libitum*. Mice were then trained to shuttle to a black barrier at the end of a linear track and back for a water reward (*alternation training*). This barrier was positioned so that there was a black background behind it (from the view of the mouse) on the wall. This track was moved to different starting positions, so the length of the track varied (**Fig. 1**). The starting position was randomly selected from 9 possible start locations for both *alternation training* and *spatial reorientation training* (spaced over a range of 56 cm – 76 cm from the center of the goal zone). Note, all calculations were actually performed in pixel coordinates. The distances listed in cm throughout the paper were estimated for visualization purposes. After reaching criterion (either asymptote, ± 6 total runs, 3 out of 4 days, or a total of 50 or more runs down the track and back), the mouse was scheduled for surgery to implant stimulating electrodes, and continued with *alternation training* every other day until the day before surgery when water deprivation ceased.

**Figure 1.**
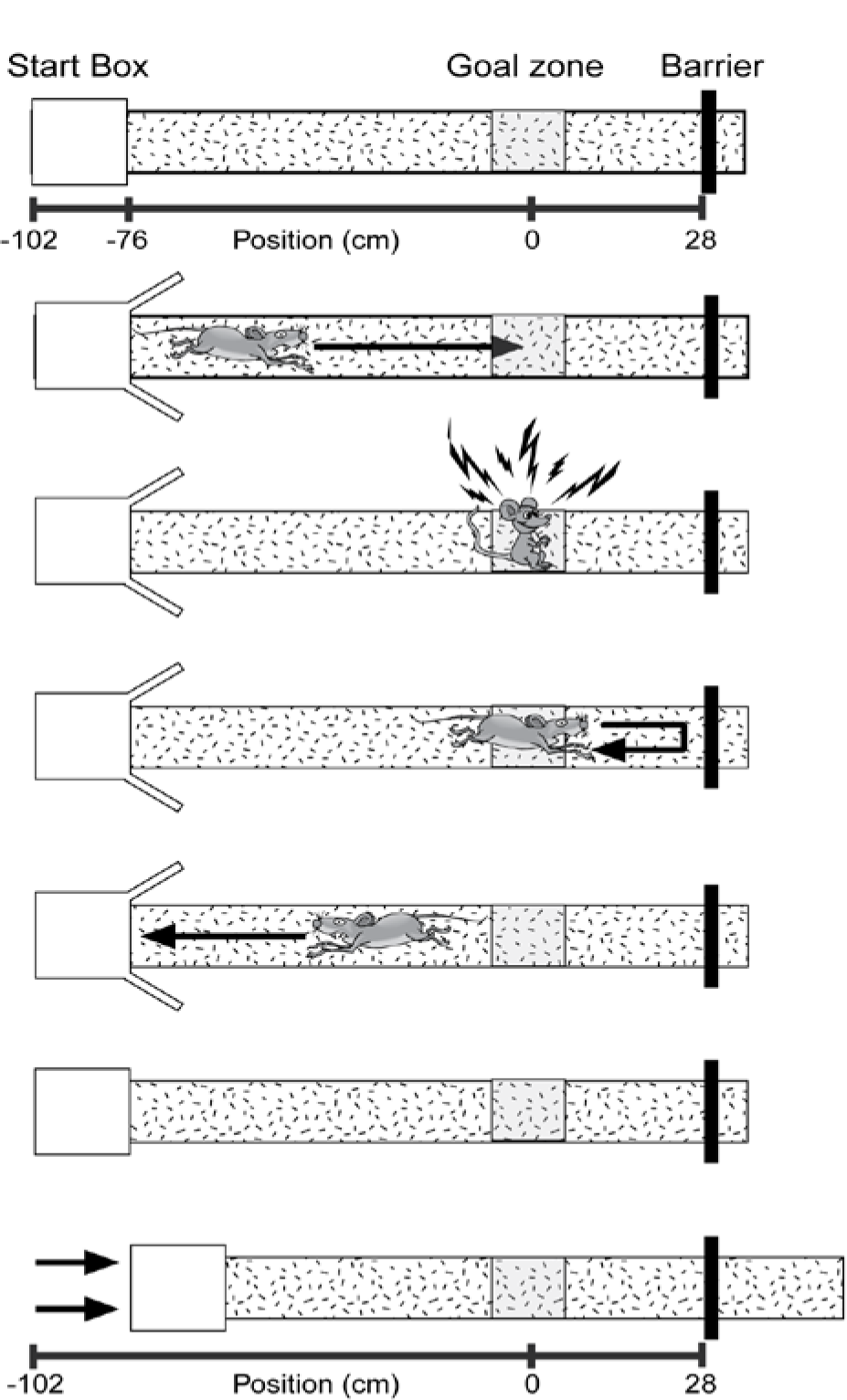
Spatial reorientation task. The goal zone (rewarded location; grey box) is always fixed within the room; however, the start box moves between trials. The sequence of events for each trial are illustrated (*top to bottom*). Each trial ends with the track being moved to a new randomly selected start location while the mouse is consuming a water reward. Thus, each trial begins with the mouse “lost” with respect to the position of the grey reward zone in the room. After leaving the start box, the mouse gets position estimates initially from self-motion (distance from the start box). As the mouse moves down the track, the position estimation is updated using room-cues. If position is successfully updated using room cues then the mouse will stop in the reward location for the required delay and obtain a brain stimulation reward. Adapted from Rosenzweig et al. (2003).

### Stimulating Electrode Implantation

Once criteria were met on the *alternation training* task, mice underwent surgery to implant bilateral stimulating electrodes targeting the left and right medial forebrain bundle (MFB; 1.9 mm posterior to bregma, 0.8 mm lateral, 4.8 mm below dura). The implantation of these stimulating electrodes allowed for intracranial stimulation of the medial forebrain bundle (Carlezon & Chartoff 2007).

### Stimulation Parameters

Following a 1-week recovery period, mice were placed in a custom 44 x 44 x 44 cm box with a circular nose poke port (Med Associates) mounted on the left half of one wall of the chamber. First, mice were shaped using manual stimulations to approach the nose poke port, then for touching the port, and finally to automatically trigger brain stimulation with beam breaks by nose poking. Following shaping, a custom MATLAB program delivered one 500 ms brain stimulation reward for each beam break in the nose poke port. Over the course of one week, settings were adjusted (171-201 Hz frequency, 50-70 μA current, electrode wire combinations) to achieve maximal response rate. No attempt was made to balance responding across genotype or age; however, we did track and compare responding (nose poke rate) across genotype and age to ensure that differences in reward strength were not likely to contribute to the observed effects. The ideal measure of stimulation effectiveness is running on a running wheel because operant conditioning is less effective in mice (Carlezon & Chartoff 2007). However, since our task requires mice to stop running for a reward, we did not want to confound the study by training mice to run for brain stimulation then training them on the reverse (stopping for brain stimulation, i.e., reversal learning). Thus, we first assessed brain stimulation responding with an operant task and this worked well for nearly all of the mice in the study. For the few mice that did not have good operant conditioning but visually appeared to respond positively to the brain stimulation reward (n = 3 mice; one 6-month 3xTg-AD female, one 6-month 3xTg-AD male, and one 6-month non-Tg male), we assessed stimulation responsiveness with a task that required alternating for stimulations at either end of the track, *stim alternation*. This was assessed after spatial re-orientation training, again to prevent a confound of reversal learning. Then, we combined responding to stimulation for these two tasks which produced a continuous distribution with all *stim alternation* values all falling below operant training values. In this combined data set we found no significant effects of genotype (3xTg-AD vs non-Tg) on response rate for either sex (ts_(8)_ < 0.37, *ps* > 0.72). Though unlikely to influence the results given that responding did not differ across genotype, it is possible that higher settings were needed to produce this similar responding in 3xTg-AD mice. Therefore, we also performed the same comparison for the two brain stimulation parameters that we adjusted: frequency and current. There were no significant differences between 3xTg-AD and non-Tg female mice for frequency (t_(8)_ = 0.65, *p* = 0.54) or current (t_(8)_ = 1.63, *p* = 0.14). For males, the list of values for 3xTg-AD and non-Tg mice were identical so statistics were performed because t = 0.

### Spatial Reorientation Training

Once mice were responding maximally, one additional refresher *alternation training* session was conducted, then *spatial reorientation training* commenced (adapted from the Rosenzweig et al 2003 task for rats). Throughout the remainder of training and testing, mice shuttled back and forth for a water reward that was delivered in the start box and consumed while the track was moved slowly and smoothly to the next randomly selected starting location (9 possible start locations spaced over a range of 56 cm – 76 cm from the center of the goal zone). Random lists were generated using Random.org (random with repeats). Next, an unmarked reward zone was added to the task (28 cm from the barrier at the end of the track) that could automatically trigger a single brain stimulation reward lasting 500 ms. This reward was only delivered if the mouse remained in this zone for a sufficient period before progressing to the end of the track (**Fig. 1**). This zone was fixed in relationship to the room and the landmarks within it and was marked in camera coordinates (i.e., there were no visible markings in the room or on the track indicating the location of the reward zone). Further, since the track was moved to new positions following each trial, any olfactory or other cues on the track could not signal the location of the reward zone. Thus, the reward zone occurred at a variety of physical locations on the track, but always in a fixed location within the room. If the mouse remained in the zone for the duration of a delay period (starting at 0.5 s), then a brain stimulation reward was delivered. The delay was fixed for a given day and was increased by 0.5 s (up to 2.5 s) each time the mouse met a criterion of similar percent correct (± 15%), 3 out of 4 days. Following completion of the linear track task, mice underwent testing on a virtual maze that was similar to the real world linear track task described above. However, analysis of that data has not yielded information that significantly extends the results presented here, thus that data is not reported here. In total training and testing for both tasks took about 2 months/mouse.

### Probe Testing

In order to ensure that mice were performing the task as intended, at the conclusion of training and testing, mice underwent two probe tests. First, to ensure that memory for each starting position was not contributing to the observed effects, after reaching criteria for the last 2.5 s delay, mice underwent an additional 2.5 s reward delay session, but with 20 possible random starting positions (same range as the random 9 condition, but with smaller increments between starting locations). Second, after mice reached asymptote on this first probe test (a drop of no more than 15% compared to asymptote from the 2.5 s reward delay), to ascertain that the mice were not using the end barrier as the sole landmark for the reward zone, a second day with 20 random starting positions was administered without the barrier at the end of the track. The same criteria as the first probe test was used for this second probe test. When piloting this behavioral protocol, we found that when the end barrier was not positioned so that it was a black barrier on a black background, performance dropped considerably during this barrier removed probe test. Therefore, care was taken to ensure the black barrier was always positioned with a black background, and a barrier removed probe test was conducted for all data reported here to ensure that no mice were using the barrier as the sole landmark for the reward location. Throughout the spatial reorientation task and probe testing, the mice were video recorded and their position was tracked using video tracking software (Neuralynx; 30 Hz frame rate). Velocity data for each position (each video frame) were extracted from the tracking data and analyzed using custom Matlab (Mathworks) scripts. We converted these velocity values to Z-scores in order to equalize performance across mice with different running speeds.

### Histology

At the conclusion of the experiment, mice were given an intraperitoneal injection of Euthasol and then transcardially perfused with 0.1 *M* phosphate-buffered saline (PBS), followed by 4% paraformaldehyde (PFA) in 0.1 *M* PBS. The whole head was post-fixed for 24 h, to allow for easy identification of the tract representing location of MFB electrodes, and then the brain was removed and post-fixed for another 24 h. Lastly, the brain was cryoprotected in 30% sucrose in 0.1 *M* PBS. Frozen sections were cut coronally with a sliding microtome at a thickness of 40 μm and split into 6 evenly spaced series.

#### 6E10

One series of sections was mounted on slides, incubated in 4% PFA for 4 min, and then rinsed with Tris-buffered saline (TBS). Next, slides were soaked in 70% formic acid for 8 – 15 min. After rinsing in TBS, slides were incubated in 0.1% Triton-X in TBS for 15 min, followed by 0.1% Triton-X and 2% bovine serum albumin (BSA) in TBS for 30 min. Sections were incubated with anti-β-amyloid 1-16 (mouse, clone 6E10, Biolegend) 1:1000 and anti-NeuN (polyclonal, rabbit, Milipore) in 0.1% Triton-X and 2% BSA in TBS for 2 days. After rinsing with TBS, slides were soaked in 0.1% Triton-X in TBS for 15 min, followed again by 0.1% Triton-X and 2% BSA in TBS for 30 min. Staining was visualized with anti-mouse-alexa-488 (1:1000) and anti-rabbit-alexa-594 (1:500) in 0.1% Triton-X and 2% BSA in TBS for 5-6 hrs. Slides were coverslipped after being rinsed with TBS. Whole slides were imaged as described below, then the coverslip was removed and DAPI (0.01 mg/ml) was added before the slides were re-coverslipped and reimaged. Except for β-Amyloid 1-16 as noted above, histology was performed on free-floating sections. Sections are permeabilized in 0.3% Triton-X and blocked in 3% Goat Serum in TBS, then incubated in primaries antibodies.

#### Thioflavin S

Anti-NeuN (1:1000), overnight, was followed by anti-rabbit-alexa-594 (1:500) for 5h. Sections were rinsed then immersed in a 1% Thioflavin S solution (Sigma) for 9 min, rinsed in dH2O, destained in 70% Ethanol for 5 min, rinsed in dH2O, and then transferred to TBS before mounting onto slides.

#### Phosphorylated tau

Anti-Phosphorylated tau (1:500, monoclonal, mouse, Thermo Scientific) with anti-NeuN (1:1000) overnight. Secondary antibodies were anti-mouse-alexa-488 (1:1000) and anti-rabbit-alexa-594 (1:500) respectively, for 6h. Sections were rinsed and mounted onto slides.

#### Parvalbumin

Sections were quench in 0.3% H_2_O_2_ in PBS for 25 minutes, then blocked in 5% goat serum in 0.5% Triton-X TBS for 90 min. Primary antibody: mouse anti-parvalbumin antibody (Sigma Aldrich) 1:2000 was added for 2 days followed by a biotinylated goat anti-mouse antibody (Sigma Aldrich) 1:500 for 90 min both in TBS with 0.5% Triton-X. Following this, A and B form the standard Vectastain ABC kit (Vector Laboratories) 1:500 in PBS was added for 1h. Staining was developed using a DAB (3,3′-Diaminobenzidine tetrahydrochloride hydrate; Sigma Aldrich) solution containing 0.05% DAB and 0.015% H_2_O_2_ in TBS. Section were rinsed in PBS and mounted onto slides. After air drying, slides were dehydrated in increasing concentration of alcohol, cleared with Hemo-De and coverslip with Fisher Chemical Permount™ Mounting Medium.

### Image Acquisition

Whole slides were scanned at 40x magnification using a scanning microscope (NanoZoomer Digital Pathology RS; Hamamatu). In addition, a subset of regions were subjectively examined in greater detail by acquiring image stacks on a confocal microscope (Olympus, FV1000).

### Region of Interest Analyses

The density of cells positive for 6e10 intracellular pathology was estimated for 3 general regions of interest (retrosplenial cortex, CA1 field of the hippocampus, and PC) which were further subdivided as follows. Dorsal CA1 (CA1d) and ventral CA1 (CA1v) were analyzed separately and CA1d was defined as all sections rostral to a point 2.55 mm posterior to bregma. CA1v was considered all sections caudal to that same point. Retrosplenial cortex was subdivided into dorsal (RSCd), ventral (RSCv), and lateral agranular (RSCagl) regions. Next, an outline was manually drawn around each region of interest (ROI) by an experimenter that was blind to group using the manual selection tool in Fiji. ROIs were based on regional boundaries provided by Allen Brain Atlas (Oh et al 2014) and cytoarchitectural differences observed in adjacent series of parvalbumin stained tissue. These boundaries are readily identifiable in NeuN-stained sections, especially when aided with adjacent parvalbumin sections, using standard cytoarchitectural criteria such as cell packing densities and thicknesses of layers. The number of labeled cells was then calculated for each cortical zone and expressed as the proportion per unit of area. Area calculations were done in pixels and converted to mm for illustrative purposes. These manual counts were acquired for a subset of the sections in the 1:6 series collected for each marker, every other section (i.e., 1:12). Manual counts were performed on 10x equivalent images extracted from 40x scans, ROI contours were drawn on subsampled 2.5x equivalent images.

### Genotyping

We received homozygous 3xTg-AD mice from Dr. Frank Laferla’s lab. We confirmed that all mice used in the experiment contained each transgene using conventional PCR. DNA was extracted from the tails of each mouse. Homozygosity was confirmed by cutting the PS1 PCR fragment with the *BstEII* restriction enzyme. Only the mutated human PS1 gene contains a *BstEII* cut site and will be cut. The absence of an uncut PCR product indicated that the mouse was indeed homozygous for the human PS1. The presence of overexpressed APP and Tau were also confirmed by PCR. The following primers were used for amplifying the PS1 transgene: 5’-CAC ACG CAA CTC TGA CAT GCA CAG GC-3’ (PS1 Forward) and 5’-AGG CAG GAA GAT CAC GTG TTC AAG TAC-3’ (PS1 Reverse). APP and Tau primers used were: 5’- GCT TGC ACC AGT TCT GGA TGG −3’ (APP Forward) and 5’- GAG GTA TTC AGT CAT GTG CT −3’ (APP Reverse) and 5’- GAG GTA TTC AGT CAT GTG CT −3’ (Tau Forward) and 5’- TTC AAA GTT CAC CTG ATA GT −3’ (Tau Reverse) respectively.

### Statistical Analyses

Data were analyzed using two-way repeated measures ANOVAs (genotype x delay, sex x delay, age x delay, and sex x ROI) followed by planned comparisons. Planned comparisons consisted of two-group *F*-tests done within the context of the overall ANOVA (Maxwell and Delaney, 2003), comparing the non-Tg group to the 3xTg-AD group. For all statistical analyses, p < 0.05 was considered significant.

### Data Availability

The datasets generated during and/or analyzed during the current study are not publicly available. However, the datasets are available from the corresponding author on reasonable request.

## Results

### Female 6-month 3xTg-AD mice are impaired at spatial reorientation

As mice learn the location of the reward zone, they slow during the approach to the zone, then stop and remain in the zone location for the required delay to obtain a brain stimulation reward (**Fig. 2A** *top*). Therefore, as a first step in assessing spatial reorientation ability in 3xTg-AD mice, we isolated velocity data (Z-scored) for one reward zone radius just before the front edge of the reward zone (track location used for data shown in **Fig. 2A** *bottom* is marked on **Fig. 2A** *top*. with an orange bar and the reward zone location is marked on **Fig. 2A** *top* with a blue bar). This measure allows for comparison of performance across reward delays (i.e., across the range of task difficulty). In addition, measuring the velocity in front of the reward zone avoids contamination from velocity change that results from running slowly through the reward zone at easier delays (e.g., 0.5s) and stopping after accidentally obtaining the reward. We compared these velocity values during the approach to the reward zone for 3xTg-AD mice to age matched non-Tg controls across reward delays (0.5 – 2.5 s). Velocity for 6-month female mice did not vary significantly across delay (**Fig. 2A**; F_(4, 32)_ = 2.25, *p* = 0.09), or across genotype (F_(1, 32)_ = 3.03, *p* = 0.12). However, there was a significant interaction between delay and genotype (F_(4, 32)_ = 3.00, *p* < 0.05). We followed up on this significant interaction with planned comparisons assessing the effect of genotype for each delay and found that 3xTg-AD female mice performed significantly worse than non-Tg age matched controls, for the 1.5 and 2 s (Fs_(1, 8)_ > 7.60, *p* < 0.05), but not 0.5, 1.0 or 2.5 s (Fs_(1, 8)_ < 3.50, *p* > 0.10) reward delays. This suggests that non-Tg mice were identifying the location of the reward zone and slowing down in preparation for stopping in the zone, but that 3xTg-AD females were not.

**Figure 2.**
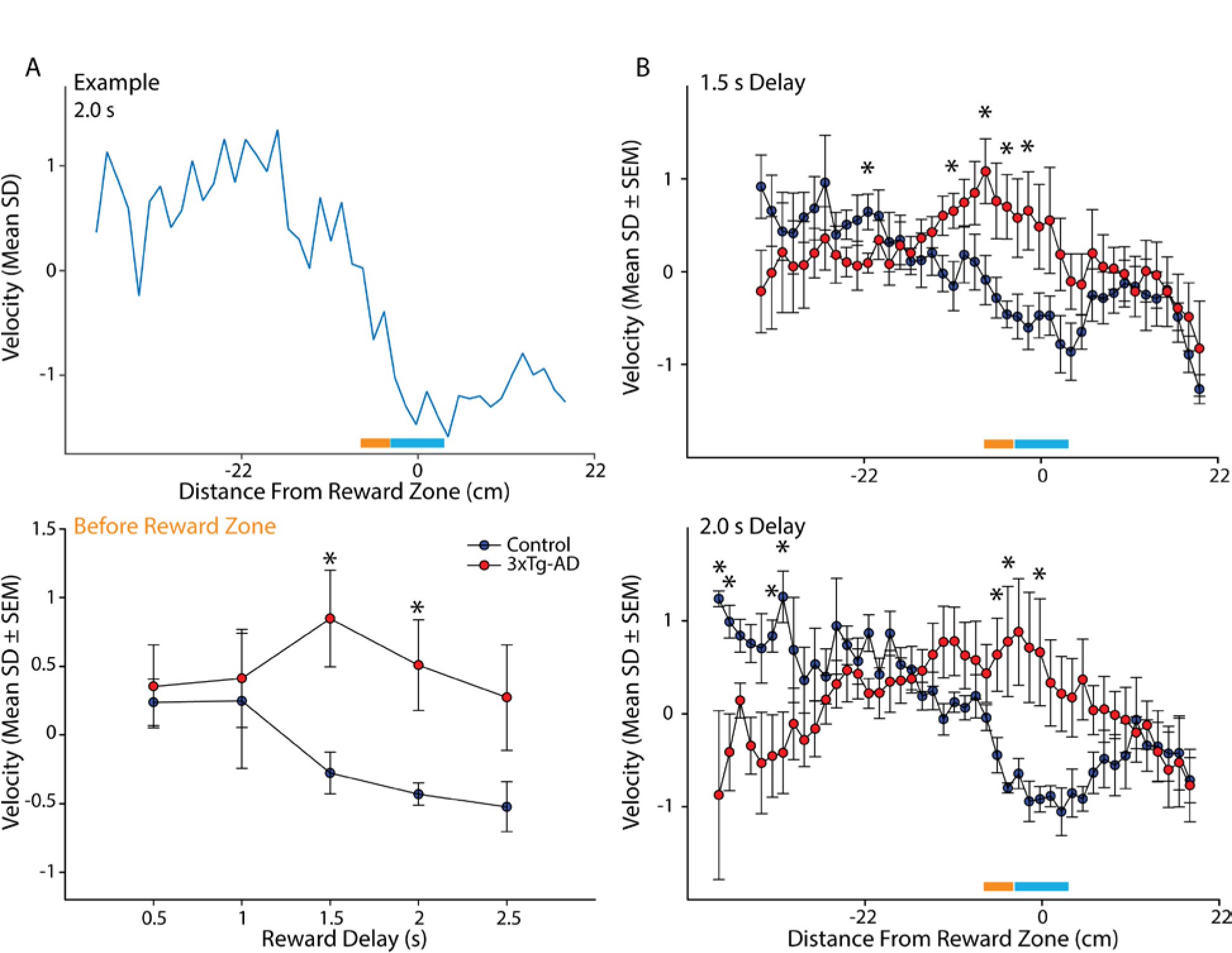
3xTg-AD female mice are impaired at the spatial reorientation task A. *Top.* Example mean Z-scored velocity for a single 6-month female non-Tg mouse for the 2.0 s reward delay. This example shows that for a longer reward delay this mouse had learned to slow as it approached the reward zone. *Bottom.* Mean (±SEM) Z-scored velocity from a reward zone radius span of track just prior to the reward zone (orange bar in **B**; reward zone is the blue bar) for each reward delay (0.5-2.5 s) for 6-month female non-Tg (blue) and 3xTg-AD (red) mice. At more difficult delays, control mice slowed down more than 3xTg-AD female mice. **B**. *Top.* Mean (±SEM) Z-scored velocity plotted along distance of the track during the *1.5 s reward delay* for 6-month female control and 3xTg-AD mice. Control mice slowed more for the approach and into the reward zone (blue bar) compared to 3xTg-AD mice. *Bottom.* Same as *Top* but for the *2.0 s reward delay*. Again, 3xTg-AD mice failed to slow in the reward zone compared to controls. \**p* < .05

Next, we examined the mean Z-scored velocities for the full length of the track individually for each reward delay where we observed a significant effect of genotype (the 1.5 and 2 s delays; **Fig. 2B**). During the *1.5 s delay*, velocity varied across the length of the track (**Fig. 2B** *top*; F_(41, 328)_ = 2.46, *p* < 0.0001), but did not vary across genotype (F_(1, 328)_ = 5.01, *p* = 0.06). However, the velocity profile varied significantly as a function of genotype and position in the room (genotype by position interaction: F_(41, 328)_ = 1.79, *p* < 0.01). Planned comparisons showed that 6-month female 3xTg-AD mice slowed less than non-Tg mice for multiple locations in the room just prior and into the reward zone (−9 cm, −5 cm, −2 cm, 1 cm; Fs_(1, 8)_ > 6.17, *p* < 0.05), but not other locations prior to the reward zone (Fs_(1, 8)_ < 5.15, *p* > 0.05), nor after the reward zone (Fs_(1, 8)_ < 3.09, *p* > 0.12). In addition, non-Tg mice ran faster than 3xTg-AD mice for 1 location near the beginning of the track (−21 cm; F_(1, 8)_ = 6.33, *p* < 0.05). Together, this pattern of data suggest that non-Tg 6-month female mice slowed more during the approach and into the reward zone than 3xTg-AD mice.

During the *2 s delay*, there was not a significant main effect of genotype on velocity (F_(1,328)_ = 0.67, *p* = 0.44), but there was a significant effect of position in the room (**Fig. 2B** *bottom*; F_(44, 328)_ = 1.73, *p* < 0.01), and a significant genotype by distance interaction (F_(44, 328)_ = 2.58, *p* < 0.0001). Planned comparisons showed that 6-month female 3xTg-AD mice slowed less than non-Tg mice for multiple locations in the room just prior and into the reward zone (−3 cm, −2 cm, 3 cm; Fs_(1, 8)_ > 6.17, *p* < 0.05). There were also four locations present towards the beginning of the track where 3xTg-AD female mice were slower than non-Tg female mice (−42 cm, −40 cm, −34 cm, −33 cm; Fs_(1, 8)_ > 5.58, *p* < 0.05). No significant difference occurred at other locations prior to the reward zone (Fs_(1, 8)_ < 4.97, *p* > 0.06), nor after the reward zone (Fs_(1, 8)_ < 2.41, *p* > 0.16). Thus, at both delays (i.e., two second highest difficulty levels), non-Tg mice slowed significantly more than 3xTg-AD mice whether measures were taken just prior to entering the reward zone or in the reward zone.

#### Probe testing suggests that female mice are using distal cues to get oriented in space

Finally, we used a probe test without the end barrier and our threshold for ‘passing’ this probe test was that a drop in the percentage of trials with successful rewards could be no more than 15%. The reference point for measuring this drop was the asymptote of the 2.5 s reward delay. Similar results were obtained for the random 20 zone list, where again, no more than a 15% drop in correct responding was observed. For both probe tests, with and without the end barrier, mice spent no more than 3 days on each in order to reach the necessary criteria.

#### Impairments in 3xTg-AD female mice are not a consequence of forgetting from day to day

Next, to look for evidence that impaired spatial reorientation may be a consequence of forgetting from day to day as reported previously for the Morris water maze (Billings et al 2005), we separated performance for the first ⅓, middle ⅓, and last ⅓ of the trials for each daily session. This was done for the mean velocity for the portion of the track just in front of the reward zone at each reward delay as described above (**Fig. 3**). If the impaired performance in 3xTg-AD mice was a consequence of impaired learning across days, there could be a significant effect observed between the last ⅓ of trials of one delay (e.g. 0.5 s) and the first ⅓ of trials of the following delay (e.g. 1.0 s). Note, this was the exact comparison used by (Billings et al 2005) to show impaired memory across days produced impaired spatial navigation. Visual inspection of the graph reveals that any difference between performance at the end of the preceding day to the beginning of the next day in 3xTg-AD mice is paralleled by a similar shift in non-Tg animals. Similarly, when this is measured statistically as in Billings et al (2005), there were no significant effects observed for either non-Tg (ts_(4)_ < 1.05, *p* > 0.28) or 3xTg-AD mice (ts_(4)_ < 1.76, *p* > 0.15). Thus, forgetting across delays was not likely to explain the impaired spatial reorientation we observed in 6-month 3xTg-AD female mice.

**Figure 3.**
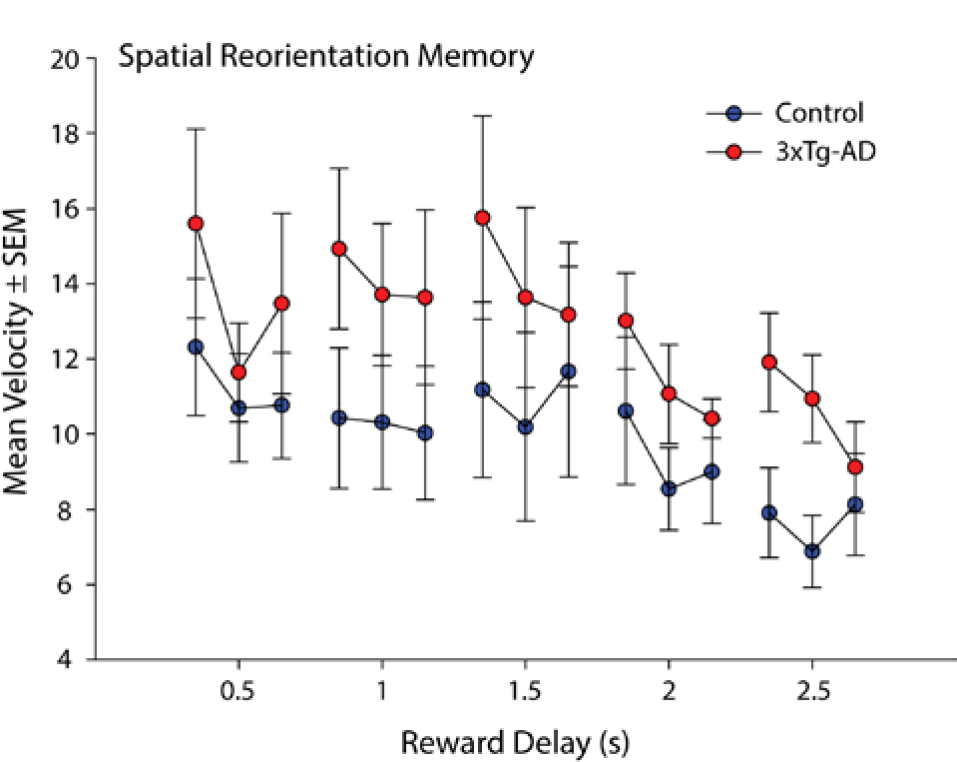
Spatial Reorientation Memory. Mean (±SEM) velocity of the first ⅓, middle ⅓, and the last ⅓ of trials from a reward zone radius span of track just prior to the reward zone for each reward delay (0.5-2.5 s) for 6-month female non-Tg (blue) and 3xTg-AD (red) mice.

### Male 6-month 3xTg-AD mice are not impaired at spatial reorientation

Next, we examined performance of 6-month male mice. As with female mice of the same age (males slightly but not significantly different in the age at first day of spatial reorientation training; t_(8)_ = 0.33, *p* = 0.75), the male mice Z-scored velocity from the portion of the track just in front of the reward zone again did not vary across delay (F_(4, 32)_ = 1.12, *p* = 0.37), nor across genotype (F_(1, 32)_ = 2.34, *p* = 0.16). However, unlike females of the same age, there was not an interaction between delay and genotype (F_(4, 32)_ = 0.52, *p* = 0.72). Thus, 6-month female, but not male, 3xTg-AD mice are impaired at the spatial orientation task.

#### Differences in markers of pathology but not average velocity corresponds to the sex difference in 6-month 3xTg-AD mice

First, we checked to see if a sex difference in non-Tg male and female mice might contribute to an apparent sex difference in 3xTg-AD mice of the same age. No significant effect of sex on velocity was present (F_(1, 32)_ = 3.89, *p* = 0.08), or a significant effect of reward delay (F_(4, 32)_ = 1.84, *p* = 0.15). In addition, there was not a significant interaction between sex and delay (F_(4, 32)_ = 1.43, *p* = 0.25). Second, to ensure that differences in running speed between the 6-month male and female mice were not contributing to the observed effects, we assessed mean raw velocities. Velocities were only included from the portion of the track where differences in Z-scored velocities were not observed (i.e., from 104 cm to 39 cm in front of the reward zone). As described above for 6-month female mice, we looked at average velocity from the portion of the track where running speed was not expected to vary in order to perform the task (i.e., baseline velocity). Baseline velocity did not vary between sex (F_(1, 16)_ = 0.70, *p* = 0.41), or genotype (F_(1, 16)_ = 0.30, *p* = 0.59), and there was not a significant interaction between genotype and sex (F_(1, 16)_ = 0.56, *p* = 0.46; Mean ±SEM velocity for non-Tg females: 10.03 ±1.95; non-Tg males: 10.16 ±0.39; 3xTg-AD females: 9.75 ±0.87; 3xTg-AD males: 11.92 ±1.65).

### Male 6-month 3xTg-AD mice have less pathology than females

When we examined the histology from these male and female mice (males slightly but not significantly different in the age at perfusion; t_(8)_ = 0.76, *p* = 0.47), it appears that amyloid beta (Aβ) pathology, as measured by the density of the subjective number of 6E10 (Aβ1-16 specific antibody) positive cells in 6-month female 3xTg-AD mice, is greater than in male mice (**Fig. 4**). The level of other markers of pathology were very low at this age (thioflavin S, phosphorylated tau; i.e., floor effect) and thus were not compared between males and females (not shown).

**Figure 4.**
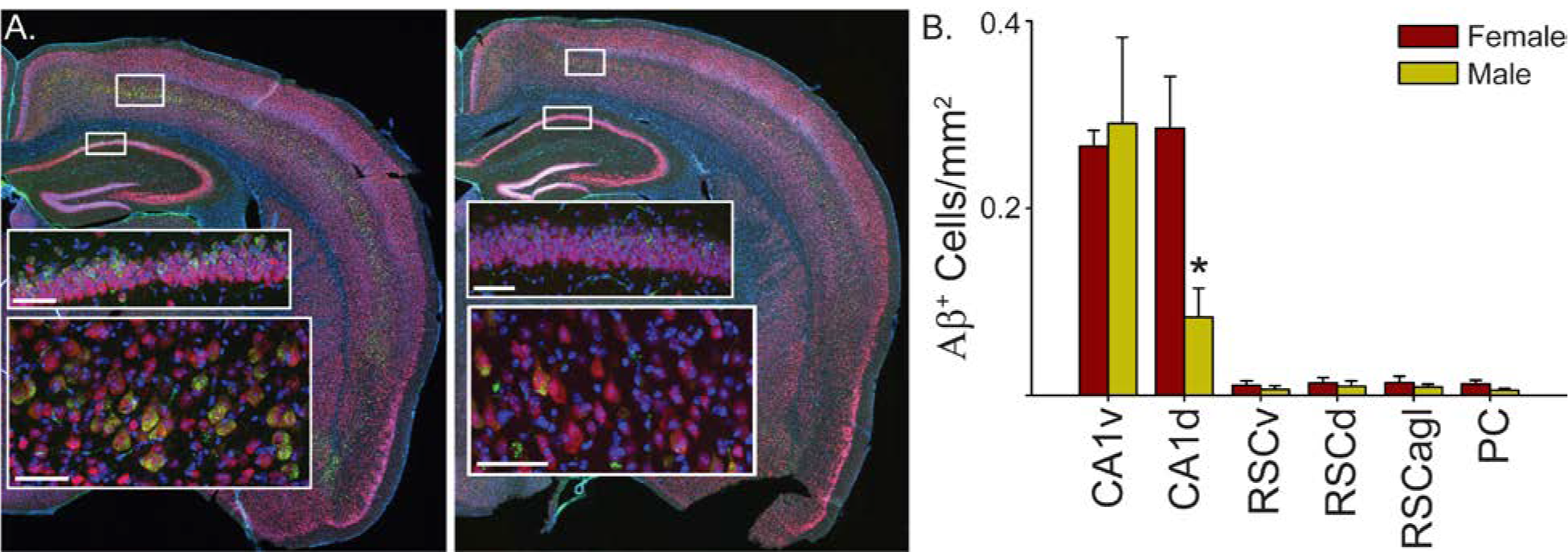
Quantitative analysis shows that 6-month 3xTg-AD female mice have significantly more 6E10 positive neurons in the dorsal hippocampus. **A**. Sections were triple labeled with DAPI (blue), NeuN (red) and 6E10 (green;-16 clone). Whole section images were acquired using a slide scanner (40x), then selected regions of interest were imaged using a confocal microscope (insets). Representative staining is shown for a 6-month 3xTg-AD female mouse (*left*) and a 6-month male mouse (*right*). At 6-months, female and male mice have only intraneuronal Aβ, and no extracellular pathology is observed. Male mice appear to have less intracellular Aβ in hippocampus and cortex. Representative confocal images are shown in the insets for cortex (*bottom inset*) and CA1 of hippocampus (*top inset*) for each hemi-section. Scale bars are 60 μm. **B.** Mean (±SEM) density of neurons positive for intraneuronal Aβ_1-16_ for 6-month male and female 3xTg-AD mice. Manual counts of 6E10 (green) positive cells were performed for the dorsal and ventral CA1 field of the hippocampus, dorsal, ventral and lateral agranular retrosplenial cortex and parietal cortex and used to calculate the mean density of 6E10 positive neurons (NeuN^+^) in each ROI for each mouse. The number of 6E10 positive cells was lower in the dorsal hippocampus (CA1d; p < 0.01) of 6-month male (gold) compared to female (maroon) mice. No significant group differences were observed for other regions. Dorsal CA1 (CA1d). Ventral CA1 (CA1v). Dorsal retrosplenial cortex (RSCd). Ventral retrosplenial cortex (RSCv). Lateral agranular retrosplenial cortex (RSCagl). Parietal cortex (PC). * p < 0.01.

Finally, for 6-month male and female 3xTg-AD mice, we quantified this subjective difference in pathology for the CA1 field of the hippocampus (that was previously linked to impaired spatial orientation in aged mice; Rosenzweig et al 2003) and two cortical regions that emerging evidence suggests may also be critical for this navigational impairment: the PC and Retrosplenial Cortex (RSC; Clark et al 2018, Hinman et al 2018). In addition, given that dorsal hippocampus is thought to be especially critical for spatial navigation, we analyzed dorsal and ventral hippocampus separately (De Hoz et al 2003, Moser et al 1993, Moser et al 1995, Pothuizen et al 2004). We found a significant effect of region (**Fig. 4B**; CA1d, CA1v, PC, RSCd & RSCv; F_(5, 40)_ = 39.26, p < 0.0001) and a significant sex by brain region interaction (F_(5, 40)_ = 4.91, p < 0.01), but not a significant effect of sex (F_(1, 40)_ = 1.98, p = 0.20). Planned comparisons indicated that female mice had significantly more pathology than male mice in dorsal CA1 (F_(1, 8)_ = 12.57, p < 0.01) but no other region (CA1v, PC, RSCd, RSCv & RSCagl; Fs_(1, 8)_ < 2.31, ps > 0.17). This suggests that a sex difference in pathology, but not sex differences in non-Tg animals or differences in running speed, may correspond to the sex difference in impaired spatial orientation in 6-month 3xTg-AD mice.

## Discussion

We found that, compared to non-Tg mice, 6-month female 3xTg-AD mice were impaired at spatial reorientation when challenged with a task requiring them use of distal cues to become reoriented. In contrast, male mice (6-months) were not impaired at the same spatial reorientation task. The impaired spatial reorientation in female mice was observed early in disease progression, when intracellular accumulation of Aβ was apparent, but prior to the appearance of extracellular pathology. The lack of impairments in male mice of the same age might correspond to a difference in the disease progression since the intracellular pathology was significantly reduced in dorsal CA1, compared to 6-month female mice. Together, our data suggest that impairments in spatial reorientation emerge very early in disease progression and that the ability to get reoriented in space may be sensitive to subtle differences in disease progression in early AD.

The impaired spatial reorientation we observed in 3xTg-AD mice is consistent with impairments in spatial learning and memory dependent tasks such as the Morris Water maze, Barnes maze, and T-maze in 3xTg-AD mice (Billings et al 2005, Cañete et al 2015, Clinton et al 2007, Davis et al 2017, Gimenez-Llort et al 2007, Nelson et al 2007, Stover et al 2015). Interestingly, the deficit in the Morris Water maze was also shown to emerge early in disease progression, before extracellular pathology was apparent; however, the spatial navigation impairment in the water maze appeared to be driven, at least largely, by forgetting from day to day with apparently normal learning across the sessions within a day (Billings et al 2005). However, we did not see evidence for such an effect in our data when making a comparison as similar as possible to Billings et al (2005). This suggests a disassociation between memory and navigation impairments at a similar point in disease progression (intracellular pathology). Thus, it might be possible to assess deficits in either memory or navigation in AD and assess the relative effects of treatments on these potentially dissociable abilities.

Previous studies showing impaired spatial navigation in mouse models of AD have assumed that mice were using distal cues to navigate; however, this was not explicitly tested (Billings et al 2005, Cañete et al 2015, Clinton et al 2007, Davis et al 2017, Gimenez-Llort et al 2007, Nelson et al 2007). Further, rats can use proximal cues to navigate the Morris Water maze (Hamilton et al 2007). Given that AD effects egocentric and allocentric navigational strategies, it is possible that navigation using proximal cues contributed at least in part to previously reported deficits. However, we explicitly tested the ability to use distal cues for navigation and found deficits early in disease progression, suggesting that use of distal cues to get oriented is dysfunctional in mouse models of amyloidosis. This failure to use distal cues suggests that future research should explore the possibility that such a deficit may explain the early dysfunction in navigating new surroundings in humans with AD (Allison et al 2016).

Interestingly, differences between males and females were only significant in the dorsal, but not ventral hippocampus, and the dorsal region is linked to performance on this and other spatial navigation tasks (De Hoz et al 2003, Moser et al 1993, Moser et al 1995, Pothuizen et al 2004, Rosenzweig et al 2003). This suggests a potential neural correlate for the early dysfunction in navigating new surroundings in humans with AD. Note, the lack of differences pathology in cortical regions does not mean that functional differences in these regions are absent. For example, Khan et al (2014) showed that pathology in entorhinal cortex and hippocampus produce a specific functional deficit in PC, even though pathology in this region was much lower than in hippocampus and entorhinal cortex.

In humans, AD has been shown to be more prevalent in females, including early stages of disease progression (Vina & Lloret 2010). Similarly, we observed that 6-month 3xTg-AD female, but not male 3xTg-AD mice, were impaired at spatial reorientation. The sex difference in spatial reorientation we reported here is consistent with previous reports showing a similar sex difference in Morris water maze deficits at a similar point in disease progression in 3xTg-AD mice (Cañete et al 2015, Clinton et al 2007). We also observed less pathology in male mice, consistent with the lack of impairment with males. Interestingly, an Aβ ELISA did not detect sex differences in pathology, suggesting the ELISA approach may be less sensitive (compared to measures of Aβ positive cells) for detecting a sex difference in pathology at least at this early point in disease progression (Clinton et al 2007). In contrast to the present results for spatial reorientation and previous reports using the water maze task where females performed worse, males were impaired on the Barnes maze spatial navigation task, but females were not (Stover et al 2015). The relationship between the sex difference and brain pathology was not described for that report, so it is unclear if variability in brain pathology or other factors contributed to this contradictory result.

The ability to successfully reorient is correlated with the realignment of the hippocampal place field map from start box centered coordinates to room centered coordinates, and aged rats have both poorer realignment to room centered coordinates and also impaired spatial reorientation (Rosenzweig et al 2003). Together, this suggests that a similar dysfunction in hippocampal place field realignment to distal cues may be the cause of the behavioral impairment observed in female 3xTg-AD mice. Mouse models of tauopathy and familial AD have shown changes in hippocampal coding and activity patterns (Cheng & Ji 2013, Witton et al 2016). Interestingly, the entorhinal cortex is also dysfunctional in AD models that mimic tauopathy, suggesting a larger brain network might be involved in spatial navigation impairments (Fu et al 2017). Future studies could directly examine the possibility that impaired hippocampal place field map alignment is ultimately responsible for the deficits we observed by recording from the hippocampus of 3xTg-AD mice while performing this task.

Navigation deficits are also a key feature of humans with AD, emerging early in disease progression (Henderson et al 1989, Weintraub et al 2012), especially when navigating in new surroundings (Allison et al 2016). The recent finding of impaired navigation in new surroundings is especially relevant for the present paper, because this is a key feature of the spatial reorientation task. This task requires the mouse to use features of the environment to determine the new starting location, which would be a critical feature of navigating new environments. This deficit, impaired navigation in new environments, is precisely the deficit observed in humans with AD. Interestingly, most human and animal studies have focused on allocentric (map-like) aspects of spatial navigation; however, a recent report showed that a distinguishing feature of human AD is the presence of both egocentric (viewer-dependent) and allocentric navigational impairments (Tu et al 2017). In fact, the spatial reorientation task requires translating between egocentric (viewer dependent) information about the landmarks and allocentric (map-like) representations of space (Byrne et al 2007, Clark et al 2018, Hinman et al 2018, McNaughton et al 1994, Oess et al 2017, Peyrache et al 2017, Whitlock et al 2012, Wilber et al 2014, Wilber et al 2017). This suggests that hippocampal and even entorhinal cortex changes may not be sufficient to explain the navigational deficits we and others have observed, and that future studies should examine both pathology profile and also function of the hippocampus and a larger brain network for spatial navigation in order to understand the changes underlying navigational impairments in AD.

In summary, we have shown a sex-dependent impairment in spatial reorientation in 3xTg-AD mice early in disease progression. The spatial reorientation task we utilized specifically taxes the use of distal cues to get oriented in space. This use of distal cues to navigate requires translation between viewer dependent and allocentric (map-like) reference frames, thus setting the stage for future studies examining the brain systems that underlie this ability in AD.

## Acknowledgments

This research was supported by grants from NIA K99 & R00 AG049090 to AAW, NIH grant AG027544 to FML, and Alzheimer’s Association NIRG-15-363477 to DBV. We thank Marina Shatskikh, Fjolla Muqolli, David Tran, Daniel Do, Christopher Sahagian, Mikko Oijala, Joshua Graham, Kayla Wellman, Andreza Melilli, Brittany Crafton, Eric Pei, and McKenzie Kirlan for technical assistance with behavioral data collection. We also thank Luis Santos-Molina and Brittany Crafton for technical assistance with image analysis.

## References

Allison SL, Fagan AM, Morris JC, Head D. 2016. Spatial Navigation in Preclinical Alzheimer’s Disease. Journal of Alzheimer’s disease: JAD 52: 77–90

Attar A, Liu T, Chan W-TC, Hayes J, Nejad M, et al. 2013. A Shortened Barnes Maze Protocol Reveals Memory Deficits at 4-Months of Age in the Triple-Transgenic Mouse Model of Alzheimer’s Disease. PLoS ONE 8: e80355

Billings LM, Oddo S, Green KN, McGaugh JL, LaFerla FM. 2005. Intraneuronal Abeta causes the onset of early Alzheimer’s disease-related cognitive deficits in transgenic mice. Neuron 45: 675–88

Braak H, Braak E. 1991. Neuropathological stageing of Alzheimer-related changes. Acta neuropathologica 82: 239–59

Byrne P, Becker S, Burgess N. 2007. Remembering the past and imagining the future: a neural model of spatial memory and imagery. Psychol Rev 114: 340–75

Cañete T, Blázquez G, Tobeña A, Giménez-Llort L, Fernández-Teruel A. 2015. Cognitive and emotional alterations in young Alzheimer’s disease (3xTg-AD) mice: Effects of neonatal handling stimulation and sexual dimorphism. Behav. Brain Res. 281: 156–71

Carlezon Jr WA, Chartoff EH. 2007. Intracranial self-stimulation (ICSS) in rodents to study the neurobiology of motivation. Nature Protocols 2: 2987

Cayzac S, Mons N, Ginguay A, Allinquant B, Jeantet Y, Cho YH. 2015. Altered hippocampal information coding and network synchrony in APP-PS1 mice. Neurobiology of Aging 36: 3200–13

Cheng J, Ji D. 2013. Rigid firing sequences undermine spatial memory codes in a neurodegenerative mouse model. Elife 2: e00647

Clark BJ, Simmons CM, Berkowitz L, Wilber AA. 2018. The retrosplenial-parietal network and reference frame coordination for spatial navigation. Retrieved from psyarxiv.com/9nb 53: 1–35

Clinton LK, Billings LM, Green KN, Caccamo A, Ngo J, et al. 2007. Age-dependent sexual dimorphism in cognition and stress response in the 3xTg-AD mice. Neurobiology of disease 28: 76–82

Davis KE, Burnett K, Gigg J. 2017. Water and T-maze protocols are equally efficient methods to assess spatial memory in 3xTg Alzheimer’s disease mice. Behav Brain Res 331: 54–66

De Hoz L, Knox J, Morris RGM. 2003. Longitudinal axis of the hippocampus: both septal and temporal poles of the hippocampus support water maze spatial learning depending on the training protocol. Hippocampus 13: 587–603

DeKosky ST, Scheff SW. 1990. Synapse loss in frontal cortex biopsies in Alzheimer’s disease: Correlation with cognitive severity. Annals of Neurology 27: 457–64

Deshmukh SS, Knierim JJ. 2013. Influence of local objects on hippocampal representations: Landmark vectors and memory. Hippocampus 23: 253–67

Flood DG, Coleman PD. 1990. Chapter 31 Chapter Hippocampal plasticity in normal aging and decreased plasticity in Alzheimer’s disease In Prog Brain Res, ed. JZ J. Storm-Mathisen, OP Ottersen, pp. 435–43: Elsevier

Forner S, Baglietto-Vargas D, Martini AC, Trujillo-Estrada L, LaFerla FM. 2017. Synaptic Impairment in Alzheimer’s Disease: A Dysregulated Symphony. Trends Neurosci 40: 347–57

Fu H, Rodriguez GA, Herman M, Emrani S, Nahmani E, et al. 2017. Tau Pathology Induces Excitatory Neuron Loss, Grid Cell Dysfunction, and Spatial Memory Deficits Reminiscent of Early Alzheimer’s Disease. Neuron 93: 533–41.e5

Gimenez-Llort L, Blazquez G, Canete T, Johansson B, Oddo S, et al. 2007. Modeling behavioral and neuronal symptoms of Alzheimer’s disease in mice: a role for intraneuronal amyloid. Neuroscience and biobehavioral reviews 31: 125–47

Hamilton DA, Akers KG, Weisend MP, Sutherland RJ. 2007. How do room and apparatus cues control navigation in the Morris water task? Evidence for distinct contributions to a movement vector. J Comp Physiol 33: 100–14

Henderson VW, Mack W, Williams B. 1989. Spatial disorientation in Alzheimer’s disease. Archives of Neurology 46: 391–94

Hinman JR, Dannenberg H, Alexander A, Hasselmo ME. 2018. Neural mechanisms of navigation involving interactions of cortical and subcortical structures. J Neurophysiol

Khan UA, Liu L, Provenzano FA, Berman DE, Profaci CP, et al. 2014. Molecular drivers and cortical spread of lateral entorhinal cortex dysfunction in preclinical Alzheimer’s disease. Nat Neurosci 17: 304–11

Liu B, Frost JL, Sun J, Fu H, Grimes S, et al. 2013. MER5101, a Novel A?1–15:DT Conjugate Vaccine, Generates a Robust Anti-Aβ Antibody Response and Attenuates Aβ Pathology and Cognitive Deficits in APPswe/PS1?E9 Transgenic Mice. J. Neurosci. 33: 7027–37

Mably AJ, Gereke BJ, Jones DT, Colgin LL. 2017. Impairments in spatial representations and rhythmic coordination of place cells in the 3xTg mouse model of Alzheimer’s disease. Hippocampus 27: 378–92

Marlatt M, Potter M, Bayer T, Praag H, Lucassen P. 2013. Prolonged Running, not Fluoxetine Treatment, Increases Neurogenesis, but does not Alter Neuropathology, in the 3xTg Mouse Model of Alzheimer’s Disease In Neurogenesis and Neural Plasticity, ed. C Belzung, P Wigmore, pp. 313–40: Springer Berlin Heidelberg

Masliah E, Mallory M, Alford M, DeTeresa R, Hansen LA, et al. 2001. Altered expression of synaptic proteins occurs early during progression of Alzheimer’s disease. Neurology 56: 127–29

McDade E, Bateman RJ. 2017. Stop Alzheimer’s before it starts. Nature 547: 153–55

McNaughton BL, Knierim JJ, Wilson MA. 1995. Vector encoding and the vestibular foundations of spatial cognition: Neurophysiological and computational mechanisms In The Cognitive Neurosciences, ed. MS Gazzaniga, pp. 585–95. Cambridge: The MIT Press

McNaughton BL, Mizumori SJY, Barnes CA, Leonard BJ, Marquis M, Green EJ. 1994. Cortical Representation of Motion during Unrestrained Spatial Navigation in the Rat. Cereb. Cortex 4: 27–39

Mesulam MM. 1999. Neuroplasticity Failure in Alzheimer’s Disease: Bridging the Gap between Plaques and Tangles. Neuron 24: 521–29

Mitchell TW, Mufson EJ, Schneider JA, Cochran EJ, Nissanov J, et al. 2002. Parahippocampal tau pathology in healthy aging, mild cognitive impairment, and early Alzheimer’s disease. Ann Neurol 51: 182–9

Moser E, Moser MB, Andersen P. 1993. Spatial learning impairment parallels the magnitude of dorsal hippocampal lesions, but is hardly present following ventral lesions. J Neurosci 13: 3916–25

Moser MB, Moser EI, Forrest E, Andersen P, Morris RG. 1995. Spatial learning with a minislab in the dorsal hippocampus. P. Natl. Acad. Sci. USA 92: 9697–701

Nelson RL, Guo Z, Halagappa VM, Pearson M, Gray AJ, et al. 2007. Prophylactic treatment with paroxetine ameliorates behavioral deficits and retards the development of amyloid and tau pathologies in 3xTg-AD mice. Experimental neurology 205: 166–76

Nitz D. 2009. Parietal cortex, navigation, and the construction of arbitrary reference frames for spatial information. Neurobiol. Learn. Mem. 91: 179–85

Nitz DA. 2006. Tracking Route Progression in the Posterior Parietal Cortex. Neuron 49: 747–56

Nitz DA. 2012. Spaces within spaces: rat parietal cortex neurons register position across three reference frames. Nat. Neurosci. 15: 1365–67

Oddo S, Caccamo A, Shepherd JD, Murphy MP, Golde TE, et al. 2003. Triple-transgenic model of Alzheimer’s disease with plaques and tangles: intracellular Abeta and synaptic dysfunction. Neuron 39: 409–21

Oess T, Krichmar JL, Röhrbein F. 2017. A Computational Model for Spatial Navigation Based on Reference Frames in the Hippocampus, Retrosplenial Cortex, and Posterior Parietal Cortex. Frontiers in Neurorobotics 11

Oh SW, Harris JA, Ng L, Winslow B, Cain N, et al. 2014. A mesoscale connectome of the mouse brain. Nature 508: 207–14

Peyrache A, Schieferstein N, Buzsáki G. 2017. Transformation of the head-direction signal into a spatial code. Nature Communications 8: 1752–52

Pothuizen HHJ, Zhang W-N, Jongen-Relo AL, Feldon J, Yee BK. 2004. Dissociation of function between the dorsal and the ventral hippocampus in spatial learning abilities of the rat: a within-subject, within-task comparison of reference and working spatial memory. European Journal of Neuroscience 19: 705–12

Rosenzweig ES, Redish AD, McNaughton BL, Barnes CA. 2003. Hippocampal map realignment and spatial learning.[see comment]. Nat. Neurosci. 6: 609–15

Stover KR, Campbell MA, Van Winssen CM, Brown RE. 2015. Early detection of cognitive deficits in the 3xTg-AD mouse model of Alzheimer’s disease. Behav. Brain Res. 289: 29–38

Tu S, Spiers HJ, Hodges JR, Piguet O, Hornberger M. 2017. Egocentric versus Allocentric Spatial Memory in Behavioral Variant Frontotemporal Dementia and Alzheimer’s Disease. J Alzheimers Dis 59: 883–92

Vina J, Lloret A. 2010. Why women have more Alzheimer’s disease than men: gender and mitochondrial toxicity of amyloid-beta peptide. J Alzheimers Dis 20 Suppl 2: S527–33

Webster SJ, Bachstetter AD, Nelson PT, Schmitt FA, Van Eldik LJ. 2014. Using mice to model Alzheimer’s dementia: an overview of the clinical disease and the preclinical behavioral changes in 10 mouse models. Frontiers in genetics 5: 88

Weintraub S, Wicklund AH, Salmon DP. 2012. The Neuropsychological Profile of Alzheimer Disease. Cold Spring Harbor Perspectives in Medicine 2

Whitlock Jonathan R, Pfuhl G, Dagslott N, Moser M-B, Moser Edvard I. 2012. Functional Split between Parietal and Entorhinal Cortices in the Rat. Neuron 73: 789–802

Wilber AA, Clark BJ, Forster TC, Tatsuno M, McNaughton BL. 2014. Interaction of Egocentric and world centered reference frames in the rat posterior parietal cortex. J. Neurosci. 34: 5431–46

Wilber AA, Skelin I, Wu W, McNaughton BL. 2017. Laminar Organization of Encoding and Memory Reactivation in the Parietal Cortex. Neuron 95: 1406–19.e5

Witton J, Staniaszek LE, Bartsch U, Randall AD, Jones MW, Brown JT. 2016. Disrupted hippocampal sharp-wave ripple-associated spike dynamics in a transgenic mouse model of dementia. J Physiol 594: 4615–30

Wood RA, Moodley KK, Lever C, Minati L, Chan D. 2016. Allocentric Spatial Memory Testing Predicts Conversion from Mild Cognitive Impairment to Dementia: An Initial Proof-of-Concept Study. Frontiers in neurology 7: 215

